# Engineering *Rhodosporidium toruloides* for sustainable production of value-added punicic acid from glucose and wood residues

**DOI:** 10.1101/2024.06.20.599976

**Authors:** Juli Wang, Dagem Zekaryas Haddis, Qiong Xiao, David C Bressler, Guanqun Chen

**Affiliations:** Department of Agricultural, Food and Nutritional Science, University of Alberta, Edmonton, Alberta, T6G 2P5, Canada

**Keywords:** *Rhodosporidium toruloides*, unusual fatty acid, punicic acid, metabolic engineering, lignocellulose

## Abstract

*Rhodosporidium toruloides* has emerged as a prominent candidate for producing single-cell oil from cost-effective feedstocks. In this study, the capability of *R. toruloides* to produce punicic acid (PuA), a representative plant unusual fatty acid, was investigated. The introduction of acyl lipid desaturase and conjugase (PgFADX) allowed *R. toruloides* to accumulate 3.7% of total fatty acids as PuA. Delta-12 acyl lipid desaturase (PgFAD2) and diacylglycerol acyltransferase 2 were shown to benefit PuA production. The strain with *PgFADX* and *PgFAD2* coexpression accumulated 12% of its lipids as PuA from glucose, which translated into a PuA titer of 451.6 mg/L in shake flask condition. Utilizing wood hydrolysate as the feedstock, this strain produced 6.4% PuA with a titer of 310 mg/L. Taken together, the results demonstrated that *R. toruloides* could serve as an ideal platform for the production of plant-derived high-value conjugated fatty acid using agricultural and forestry waste as feedstock.

## 1. Introduction

Punicic acid (PuA; 18:3Δ^9*cis*, 11*trans*, 13*cis*^) is a plant-derived conjugated fatty acid with various health-promoting benefits (Holic et al., 2018). In addition, due to the presence of conjugated double bonds, conjugated fatty acids have strong chemical reactivity and polymerization ability, making them useful as cross-linking agents in oleochemical industries (Adekunle, 2015; He et al., 2014). The primary natural source of PuA is pomegranate (*Punica granatum*) seed oil, which contains up to 80% of its fatty acids as PuA (Khoddami et al., 2014; Paul and Radhakrishnan, 2020).

In developing pomegranate seeds, oleic acid generated via the *de novo* fatty acid biosynthesis is esterified to phosphatidylcholine (PC). Pomegranate delta-12 acyl lipid desaturase (PgFAD2) then catalyzes the desaturation of oleic acid at the *sn-2* position of PC, resulting in linoleic acid (LA, C18:2). A bifunctional fatty acid desaturase and conjugase PgFADX then further converts LA to PuA on PC (Iwabuchi et al., 2003). PuA can be released into the cytosol in the form of acyl-CoA and then channelled to the storage lipid triacylglycerol (TAG) (Fig. 1A). Various acyltransferases play important roles in the acyl-editing and TAG assembly process for the efficient accumulation of unusual fatty acids (Cahoon and Li-Beisson, 2020). For instance, diacylglycerol acyltransferase (DGAT) catalyzes the committed last step in TAG biosynthesis and certain DGAT2s were shown to have a substrate preference for unusual fatty acids (Chen et al., 2022; Shockey et al., 2006). In addition, phosphatidylcholine: diacylglycerol cholinephosphotransferase (PDCT) catalyzes the interchange of diacylglycerol and PC and facilitates the enrichment of unusual fatty acids in TAG (Demski et al., 2022).

**Fig. 1.**
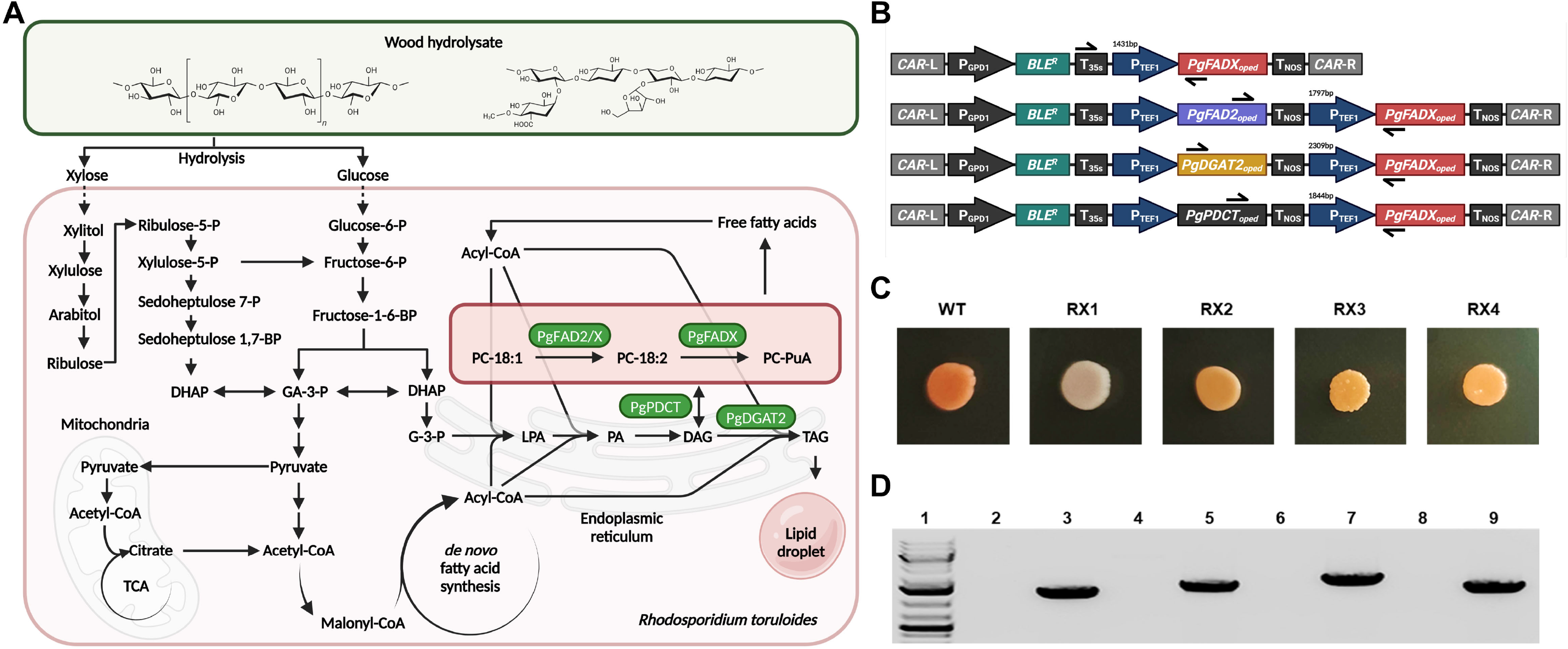
Metabolic engineering of PuA-producing *R. toruloides*. A) Illustration of the metabolic pathway involved in PuA biosynthesis in *R. toruloides*. Abbreviations: PgDGAT2, *P. granatum* acyl-CoA: diacylglycerol acyltransferase 2; PgFAD2, *P. granatum* fatty acid desaturase 2; PgFADX, *P. granatum* fatty acid desaturase and conjugase; PgPDCT, *P. granatum* phosphatidylcholine: diacylglycerol cholinephosphotransferase; GA-3-P, glyceraldehyde 3-phosphate; DHAP, dihydroxyacetone phosphate; G-3-P, glycerol-3-phosphate; LPA, lysophosphatidic acid; PA, phosphatidic acid; DAG, diacylglycerol; TAG, triacylglycerol; PC, phosphatidylcholine. B) Donor DNA structures and genomic integrations of codon-optimized *PgFADX, PgFAD2, PgDGAT2,* and *PgPDCT* into *R. toruloides.* Pathway and donor DNA figures were created with Biorender. C) Colony colour of *R. toruloides* wild-type strain as well as engineered strains RX1, RX2, RX3 and RX4. D) Detection of positive transformants. Lane 1, GeneRuler 1 kb Plus DNA ladder; Lane 2/4/6/8, *R. toruloides* wild type strain; Lane 3/5/7/9, detection of integrated cassettes in strain RX1, RX2, RX3 and RX4, respectively.

Despite the high concentration of PuA in pomegranate seed oil, several challenges remain with the production of this unusual fatty acid via traditional agriculture. For instance, the PuA content in different pomegranate cultivars varied substantially (Fadavi et al., 2006), and the oil yield from pomegranate seeds is lower than traditional oilseed crops, such as canola (Rahman et al., 2013). Although a few proof-of-concept studies have produced PuA in plants such as the model plant *Arabidopsis thaliana* and oilseed crop *Brassica napus* under laboratory conditions (Mietkiewska et al., 2014; Xu et al., 2020), more efforts are needed to develop elite crop cultivars for cost-effective commercial production of PuA. In contrast, microbial production via fermentation provides a sustainable, effective, and scalable approach with significant benefits of improved yields, cost-effectiveness, and shorter production cycles, and thus is considered a promising alternative for the production of plant-derived unusual fatty acids. In a previous study, baker’s yeast (*Saccharomyces cerevisiae*) was modified to produce 3.7% of total fatty acids as PuA using LA precursor feeding (Wang et al., 2021). In another study, the metabolic engineering of *Schizosaccharomyces pombe*, a yeast with high oleic acid content, led to 25.1% of total fatty acid as PuA (Garaiova et al., 2017). However, since *S. cerevisiae* and *S. pombe* have very limited ability to accumulate lipids, the PuA titer was only 7.2 mg/L and 38.7 mg/L, respectively. In this regard, it is attractive to search and develop new microbial platforms for the production of this plant-derived unusual fatty acid via metabolic engineering.

Some yeast species are well-recognized for their potential to produce single-cell oil. Oleaginous yeasts, such as *Yarrowia lipolytica* and *Rhodosporidium toruloides*, can accumulate more than 20% of their biomass as lipids. For instance, *Y. lipolytica* can grow on various carbon sources including fatty acids, glucose, fructose, or glycerol, and accumulate 36% of its biomass as lipids (Zhang et al., 2014), and it has been tested for the production of several unusual fatty acids (Park and Hahn, 2024; Urbanikova et al., 2023; Xue et al., 2013). *R. toruloides*, a non-conventional yeast capable of concomitant synthesis of lipids and carotenoids, is typically regarded as a promising candidate for biofuel production due to its excellent lipid accumulation capability, which often exceeds 60% of dry cell weight (González-García et al., 2017). More interestingly, *R. toruloides* has the natural ability to use a wide range of substrates, including pentose sugars such as xylose (Adamczyk et al., 2023; Coradetti et al., 2023). This advantage makes *R. toruloides* promising for sustainable bioindustry and agriculture applications since xylose assimilation capability is essential for converting lignocellulosic biomass, the most abundant raw material derived from agriculture and forestry waste, into value-added unusual fatty acids.

In this study, we evaluated the potential of engineering the nonconventional yeast *R. toruloides* for the production of PuA, a plant-derived value-added unusual fatty acid. *R. toruloides* transformed with *PgFADX* was able to accumulate PuA to 3.7% of total fatty acids. By further integrating codon-optimized *PgFAD2* or *PgDGAT2* into *R. toruloides’* genome, PuA contents were significantly improved. The best engineered strain *R. toruloides* RX2 containing both *PgFADX* and *PgFAD2* accumulated 12% of total lipid as PuA using glucose as substrate and a titer of 451.6 mg/L PuA was achieved in the flask cultivation. Lipid fraction analysis revealed that, when cultured under the nitrogen-limited condition, the recombinant *R. toruloides* cells accumulated a higher relative content of PuA in TAG than in polar lipid (PL). Finally, we tested if *R. toruloides* RX2 could use wood hydrolysate as the feedstock for PuA production, and the results revealed that the strain could accumulate 6.4% of its lipid as PuA, which demonstrated a good potential of *R. toruloides* in converting low-value agricultural waste into value-added PuA.

## 2. Materials and Methods

### 2.1 Strains, plasmids and culture conditions

Strains and plasmids used in this study are listed in Table 1 and Table S1. *Escherichia coli* DH5α was used for routine plasmid construction and preparation. The wild-type *R. toruloides* ATCC 204091 (formerly known as *Rhodotorula glutinis*) was obtained from the American Type Culture Collection (ATCC, Manassas, VA, USA). To construct mutants with various gene deletions, the 1 kb upstream and downstream flanking sequences of the *CAR2* gene were amplified from the *R. toruloides* genomic DNA and linked with expression cassettes containing the zeocin-resistance gene and the target gene. *CAR2* encodes lycopene cyclase, which is responsible for carotenoid pigment biosynthesis (Qi et al., 2020). Insertion into the *CAR2* gene by homologous recombination leads to white colonies due to the disruption of carotenoid biosynthesis, which serves as a visual marker besides antibiotic selection for identifying transformants with successful integration. *EcoRV* restriction sites were introduced to both ends of the flanking sequences in the primer design. The zeocin-resistance gene was under the control of *GPD1* promoter and *35S* terminator. The target genes, including codon-optimized *PgFADX*, *PgFAD2*, *PgDGAT2*, and *PgPDCT*, were placed under the control of the *translation elongation factor 1* (*TEF1*) promoter, which is a strong constitutive promoter with robust performance under various growth conditions and *NOS* terminator (Nora et al., 2019) (Fig. 1B). To release donor DNAs from the plasmid backbone, either PCR or double digestion of *EcoRV* sites flanking the donor DNA was conducted.

**Table 1.**
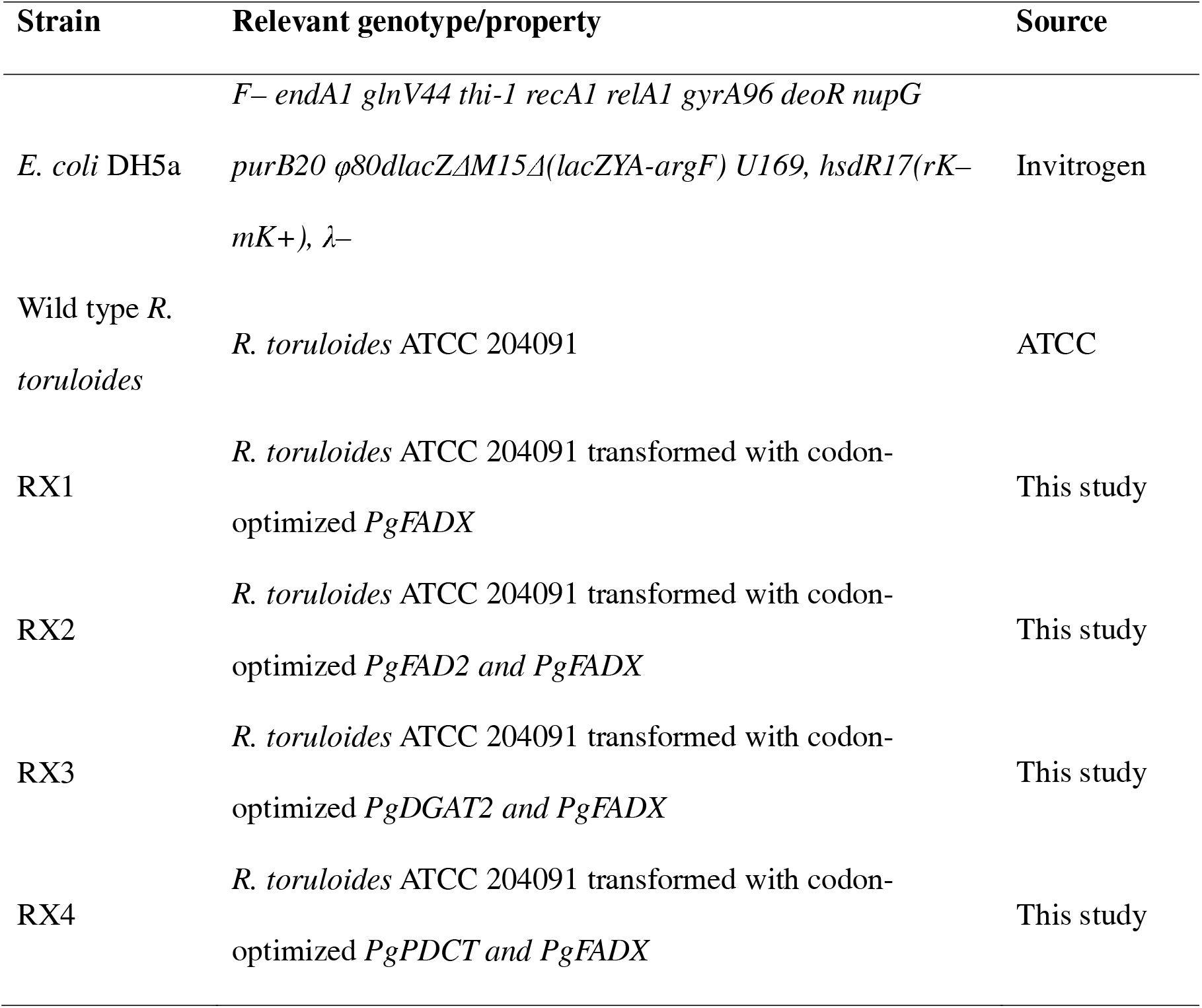
Strains used in this study.

*E. coli* was cultured in Luria-Bertani (LB) medium at 37 °C with constant shaking at 200 rpm. Kanamycin (50 μg/L) was added to maintain a stable inheritance of plasmids. *R. toruloides* was routinely maintained with yeast extract peptone dextrose (YPD) medium containing 10 g/L yeast extract, 20 g/L peptone, and 20 g/L glucose at 30 °C. To induce lipid production, seed culture of *R. toruloides* was first grown in a 50 mL falcon tube containing 5mL YPD medium at 30 °C and 250 rpm and then inoculated into 1 L shake flasks containing 200 mL of nutrient-limited media (100 g/L glucose; 0.1 g/L NaNO_3_, 4.5 g/L KH_2_PO_4_, 0.2 g/L MgSO_4_·7H_2_O, and 0.11 g/L CaCl_2_·2H_2_O) to an initial OD_600_ of 0.8 (González-García et al., 2017).

### 2.2 RNA extraction, cDNA synthesis and gene expression analysis

Total RNA was extracted from various recombinant *R. toruloides* strains cultured in the nitrogen-limited medium using the Spectrum Total RNA Kit (Sigma-Aldrich, Oakville, Canada). First-strand cDNA was synthesized with the SuperScript IV first-strand cDNA synthesis kit (Invitrogen, Carlsbad, USA), according to the manufacturer’s instructions, and then diluted 10 times and used as the template for the analysis of *PgFADX* expression with quantitative RT-PCR (qPCR). *R. toruloides GPD1* was used as the internal reference. qPCR was conducted with three biological replicates on a StepOnePlus Real-Time PCR System (Applied Biosystems, USA) using the GB-Amp™ Sybr Green qPCR Mix (GeneBio, Burlington, Canada) according to the manufacturer’s instructions. Results were analyzed using the comparative Ct method (2^-ΔΔCt^ method).

### 2.3 Chemical transformation of *R. toruloides*

The yeast transformation was performed using the lithium acetate and PEG method (Nora et al., 2019; Tsai et al., 2017). Briefly, the wild-type *R. toruloides* strain was first cultured in YPD medium at 30°C and 200 rpm overnight. The next day, the cell was diluted with fresh medium to OD_600_ of 0.2 and grown to OD_600_ of 0.8. Biomass was then recovered, washed and resuspended in 1 mL 100 mM lithium acetate. After centrifugation, 10 μL single-stranded salmon sperm DNA (10 mg/mL), 240 μL PEG4000, 36μL lithium acetate (1.0 M), and 5 μg DNA of interest was added to the cells, which were then incubated at 30 °C for 40min. Subsequently, 34 μL dimethylsulfoxide was added to the mixture, followed by heat shock at 42 °C for 15 min. Transformed yeast cells were allowed to recover overnight at 30 °C in the YPD medium with shaking. The cells were plated on YPD agar plates containing 150 μg/mL of Zeocin and incubated at 30 °C.

### 2.4 Preparation of wood hydrolysate

Enzymatic hydrolysis of a wood pulp composed of 79.1% cellulose, 21.2% hemicellulose and 4.0% lignin (Northern Bleached Hardwood Kraft (NBHK)) was conducted at a solid concentration of 10 % (w/v) in 50 mM sodium acetate buffer (pH 4.8) and 20 FPU/g of cellulase enzyme at 50 °C for 24 h with 200 rpm agitation. The liquid enzyme hydrolysate was analyzed for glucose and xylose sugar yields on an Agilent 1200 HPLC coupled with a Refractive Index Detector (Agilent 1100, Santa Clara, CA, USA). Sugars were separated on an HPX-87h column (Bio-Rad Aminex, Hercules, CA, USA) with 20 µL injection volume and 50 mM of sulfuric acid as the mobile phase at a flow rate of 0.5 mL/min at 60 °C for 40 min.

### 2.5 Lipid extraction

Yeast biomass was harvested from liquid culture via centrifugation and then mixed with 800 µL of a precooled lipid extraction mixture containing chloroform and isopropanol (2:1, v/v) along with glass beads (0.5mm). Butylated hydroxytoluene (BHT) was added as the antioxidant at a final concentration of 0.01% to protect PuA from degradation. Subsequently, *R. toruloides* cells were disrupted via three cycles of bead beating (1-minute duration each) using a Biospec bead beater (Biospec, Bartlesville, OK), with a 2-minute cooling on ice between each cycle. The extraction process was repeated twice. The collected organic phase was pooled together, dried under nitrogen, resuspended in chloroform and stored under - 20 °C until further analysis.

### 2.6 Separation of lipid class using thin-layer chromatography

For the detailed analysis of lipid classes, extracted lipids were separated using silica gel-coated thin-layer chromatography (TLC) (0.25 mm Silica gel, DCFertigplatten, Macherey-Nagel, Germany) in a solvent mixture consisting of hexane, diethyl ether, and acetic acid in a 70:30:1 volume ratio (Mietkiewska et al., 2014). Once the TLC plates were developed, lipid fractions were visualized using a 0.05% primulin dissolved in acetone: water (8:2 v/v), which allows for the non-destructive identification of lipid bands. Following the identification of lipid fractions, the bands corresponding to the target lipid classes were scraped from the plates. These collected fractions were then subjected to lipid extraction and base-catalyzed lipid derivatization.

### 2.7 Positional analysis of triacylglycerol with enzymatic hydrolysis

The distribution of fatty acids within the *sn-2* and *sn-1/3* positions of TAG was conducted with enzymatic hydrolysis (Luddy et al., 1964; Xu et al., 2020). Briefly, TAG fractions recovered from the TLC plate were moved to clean tubes and dried under a stream of nitrogen. For enzymatic hydrolysis, lipids were mixed with 1 mL Tris-HCl buffer (1 mM, pH 8.0), 100 μL 2.2% CaCl_2_, and 250 μL 0.1% deoxycholate. This mixture was then vortexed and sonicated briefly for lipid emulsification. After being pre-warmed for 30 seconds at 40°C, 20 mg of pancreatic lipase (pancreatic lipase type II, Sigma) was added to the mixture to initiate hydrolysis. This enzyme specifically targets the acyl chain on the *sn-1/3* positions in TAGs. The reaction was continued for 3 minutes at 40°C and quenched with 500 μL of 6 M HCl. Lipids were then extracted twice with diethyl ether, followed by a secondary TLC separation. The band corresponding to the *sn-2* monoacylglycerol fraction was recovered, transmethylated and analyzed on the gas chromatograph (Xu et al., 2020).

### 2.8 Lipid transmethylation and analysis

Briefly, for preparing fatty acid methyl esters (FAMEs), transmethylation was conducted via a base-catalyzed method using 1 mL of 5% sodium methoxide dissolved in methanol (Mietkiewska et al., 2014). After incubation in the dark at 30 °C for 1 hour, the transmethylation reaction was stopped by adding 1.5 mL of 0.9% (w/v) sodium chloride solution. FAME was extracted twice with hexane, dried under nitrogen, and analyzed on an Agilent 6890N Gas Chromatograph equipped with a 5975 inert XL Mass Selective Detector (for qualification) and a Flame Ionization Detector (for quantification). FAMEs were separated on a capillary DB23 column (30 222 m×0.25 mm×0.25 µm, Agilent Technologies, Wilmington, DE, USA) with split injection (5:1 split ratio, 1 µL injection) using the following program: 165 °C for 4 min, increased to 180 °C (10 °C/ min) and held for 5 min, and then increased to 230 °C at 10 °C/min and held for 5 min.

## 3. Results and Discussion

### 3.1 Reconstitution of PuA synthesis in R. toruloides via the heterologous expression of PgFADX

Since controlling the carbon/nitrogen ratio in the culture medium significantly impacts lipid metabolism in oleaginous microorganisms, to improve the PuA production and lipid content, the wild-type and recombinant *R. toruloides* were cultured under the nitrogen-limited medium optimized for lipid induction (González-García et al., 2017). As shown in Fig. 2A, after 96 hours of cultivation under nitrogen-limited conditions, the wild-type yeast cell accumulated C18:1 and C18:2 to 47% and 10% of total fatty acids, respectively. In contrast, there’s only a limited amount of C18:3 fatty acids (<2.7%) and no conjugated fatty acid was detected in the wild-type strain. The combined C18 fatty acid occupied over 70% of total fatty acids, which is conducive to PuA accumulation since C18:1 and C18:2 are the important fatty acid precursors to PuA synthesis.

**Fig. 2.**
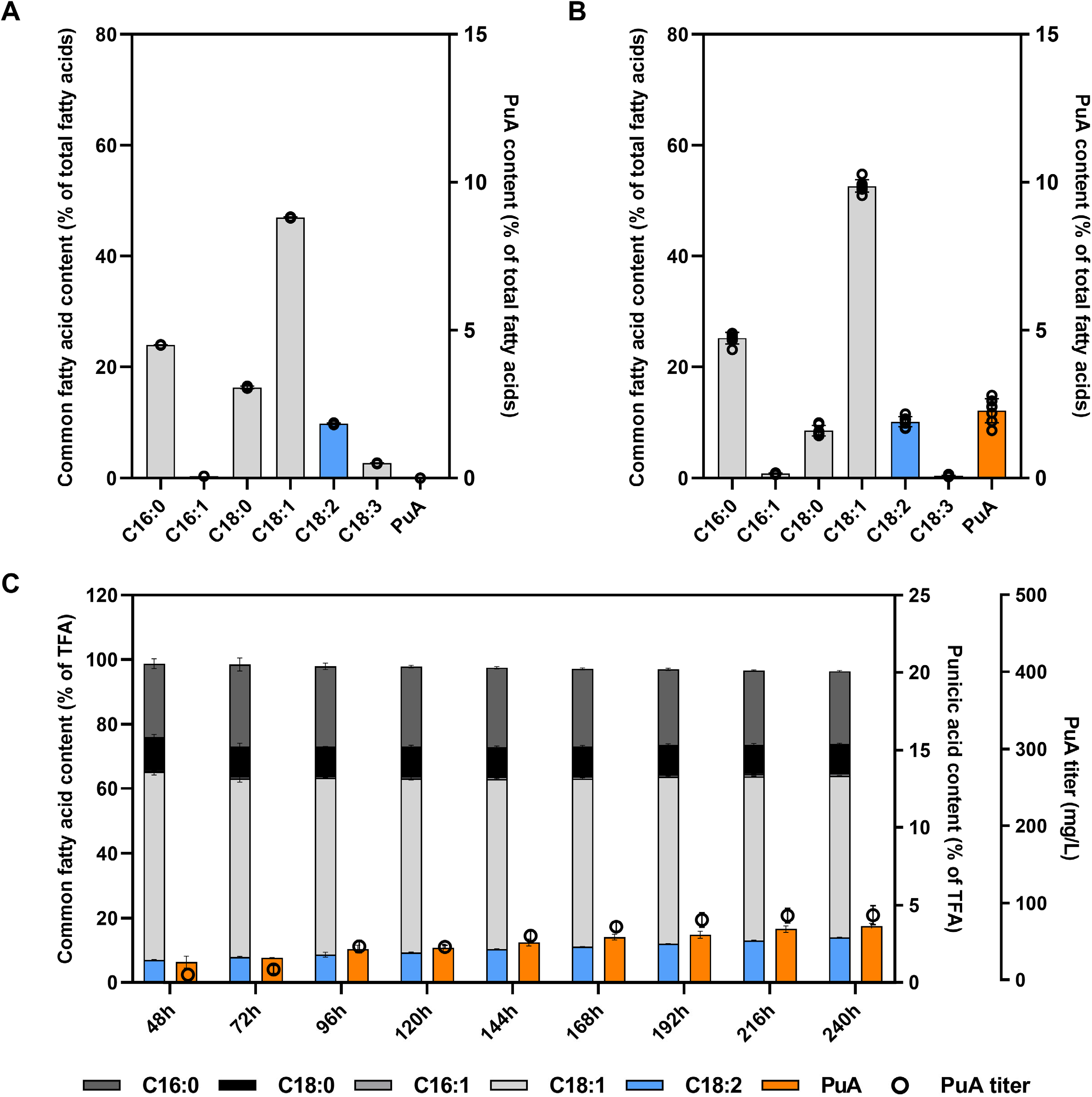
Genomic integration of the pomegranate *PgFADX* expression cassette led to PuA-producing *R. toruloides* strain RX1. A) Fatty acid composition of wild-type *R. toruloides* cultured in chemically defined medium. B) Integration of *PgFADX* led to PuA product. C) Fatty acid composition and PuA titer of RX1 strain from 48h to 240h. Data represent mean ± SD of triplicates.

For the reconstitution of the PuA synthetic pathway in *R. toruloides*, a donor DNA cassette was first assembled (Fig.1B). Since *R. toruloides* genes have very high (∼60%) GC contents (Hu and Ji, 2016), the gene encoding pomegranate-derived *PgFADX* was codon optimized to accommodate *R. toruloides*’ specific codon usage bias (Table S2) and integrated into *R. toruloides*’ genome targeting the *CAR2* region, which was showed to have higher gene editing efficiency compared to other commonly used loci, such as *URA3* (Otoupal et al., 2019). As shown in Fig. 1C, after the transformation of *PgFADX*, strains with white colonies were obtained on the zeocin agar plate. Subsequent analysis detected the *PgFADX* cassette on the genomic DNA recovered from positive transformants (Fig. 1D).

In order to examine their PuA-accumulating ability, seven single white colonies were tested in the nitrogen-limited medium. As shown in Fig. 2B, the transformation of *PgFADX* led to 1.6%-2.6% of total fatty acids as PuA, with an average content of 2.3%. Notably, the linolenic acid content in transformed *R. toruloides* decreased to less than 0.3% (Fig. 2B). Since linolenic acid synthesis competes with PuA synthesis for LA substrate, in a previous study with transgenic plant, a mutant *Arabidopsis* line lacking fatty acid desaturase 3 activity was used to limit the competition and improve PuA production (Mietkiewska et al., 2014). Although a bifunctional Δ12/Δ15 fatty acid desaturase was suggested to carry out further desaturation converting LA into linolenic acid in oleaginous yeast such as *R. toruloides* and *Lipomyces starkeyi* (Y. Liu et al., 2021; Matsuzawa et al., 2018), the very low content of linolenic acid content in the transgenic *R. toruloides* strains indicated that PgFADX may effectively compete with *R. toruloides*’ endogenous desaturase for LA substrate. Therefore, removing the activities of native bifunctional desaturases may not be necessary.

To further characterize the influence of *PgFADX* overexpression, the transformant with the highest PuA content, designated as RX1, was tested in a 10-day shake flask cultivation. As shown in Fig. 2C, the level of LA and PuA in strain RX1 gradually increased in the 10-day period, and PuA accounted for 3.7% of total fatty acids at the end of cultivation. A concomitant increase in LA and PuA was observed, which was possibly due to PgFADX’s bifunctional activity to produce both LA and PuA (Garaiova et al., 2017). On the contrary, the content of oleic acid level was reduced from 58% to 50% between day 2 and day 10. The PuA titer at the end of the cultivation reached 88.8 mg/L, which was higher than the level previously achieved in *S. cerevisiae* and *Y. lipolytica* with single *PgFADX* expression (Urbanikova et al., 2023; Wang et al., 2021). Taken together, the results indicated that *R. toruloides* has great potential to become a yeast platform for producing plant-derived PuA.

### 3.2 Coexpression of *PgFADX* with *PgFAD2* or *PgDGAT2* significantly improved PuA content in recombinant *R. toruloides*

Considering the importance of LA precursor to PuA synthesis, we further constructed the *PgFAD2* and *PgFADX* coexpression cassette, aiming to provide more LA precursor to PgFADX on top of *R. toruloides*’ native LA synthesis. Moreover, since the channelling of unusual fatty acids in the natural producer also requires a series of downstream enzymes such as DGAT2 and PDCT (Demski et al., 2022; Shockey et al., 2006), strains coexpressing *PgFADX* with *PgDGAT2* or *PgPDCT* were also constructed, respectively (Table 1). Only a few transformants harboring the coexpression of two genes were recovered from the zeocin selection plate, and all of them displayed an orange pigmentation (Fig.1C), indicating the incomplete *CAR2*-targeted insertion and disruption by the cassettes. The incomplete disruption may caused by the much longer length of the co-expression cassettes, in comparison to the single *PgFADX* expression cassette (Fig.1B), or by the preference for nonhomologous end joining (NHEJ) over homologous recombination (HR) for DNA repair in the *R. toruloides* strain (Schultz et al., 2019). Indeed, earlier studies indicated that high *CAR2* deletion efficiency up to 75.3% was only obtained in the KU70-deficient *R. toruloides* strain, which has a mutation of Ku70/80 regulatory DNA-binding subunits that is critical to the NHEJ system, whereas in the wild-type *R. toruloides* strain, the targeted deletion frequency of *CAR2* was only 10.5% (Koh et al., 2014). The donor DNA cassettes generated for coexpression appear to be randomly integrated into the *R. toruloides* genome by NHEJ, in line with a previous study using a similar transformation method (Tsai et al., 2017).

To take into account the position effect, in which surrounding genetic elements may enhance or inhibit gene expression, different transformants were examined for their varying abilities to accumulate PuA. As shown in Fig. 3A, *PgFADX*-*PgFAD2* coexpression led to 5.8%-9.6% of total fatty acids as PuA, with an average of 7.4%, representing a 2.3-fold increase compared to *PgFADX* single expression (Fig. 2). In addition, the average LA content increased by 30% at the expense of oleic acid content (Fig. 3A). Similarly, the coexpression of *PgDGAT2* with *PgFADX* resulted in a 2-fold rise in PuA content, whereas LA content was decreased by 14% (Fig.3B). In contrast, the coexpression of *PgPDCT* with *PgFADX* only increased the PuA content by merely 17% (Fig. 3C). The transformants with highest PuA content for *PgFADX*-*PgFAD2*, *PgFADX*-*PgDGAT2*, and *PgFADX*-*PgPDCT* coexpression, designated as strains RX2, RX3, and RX4, were subsequently investigated in shake flask culture (Fig. 3D, 3E and 3F). By the end of the 10-day cultivation, LA and PuA accounted for 13.8% and 12% of total fatty acid in RX2, respectively (Fig. 3D). The final lipid content was 66% of dry cell weight, and PuA titer of strain RX2 reached 451.6 mg/L, which was 5-fold that of RX1. In strain RX3, the PuA levels reached 8% of total fatty acids at the end of cultivation and led to a final PuA titer of 285.4 mg/L (Fig. 3E), which was 3.2-fold that of RX1.

**Fig. 3.**
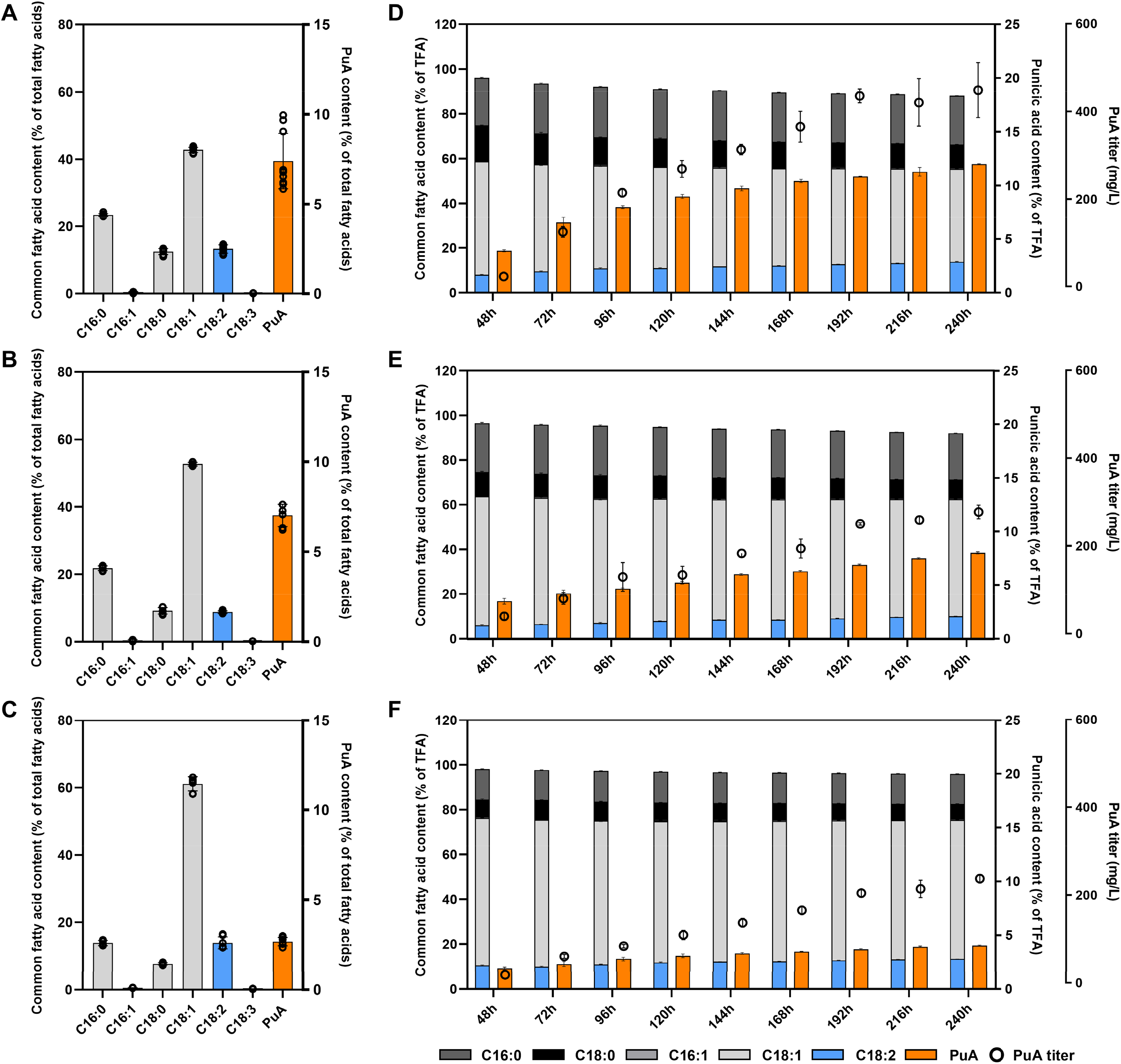
Coexpression of pomegranate *PgFAD2*, *PgDGAT2*, and *PgPDCT* respectively with *PgFADX* led to increased PuA production. A-C) Fatty acid composition of *R. toruloides* strains coexpressing *PgFAD2-PgFADX, PgDGAT2-PgFADX, and PgPDCT-PgFADX*. D-F) Fatty acid composition and PuA titer of *R. toruloides* strains RX2, RX3 and RX4 from 48h to 240h. Data represent mean ± SD of triplicates.

To determine whether the increase in PuA production resulted from changes in *PgFADX* expression levels or from the beneficial effects of the coexpressed enzymes, the relative expression levels of *PgFADX* in RX2, RX3, and RX4 were also quantified using RT-PCR (Fig. 4A). Results showed *PgFADX* expressions were not higher in RX2, RX3, and RX4 when compared to the level in strain RX1. Collectively, the results indicated that the introduction of *PgFAD2* or *PgDGAT2* could significantly further improve PuA production in *R. toruloides*.

**Fig. 4.**
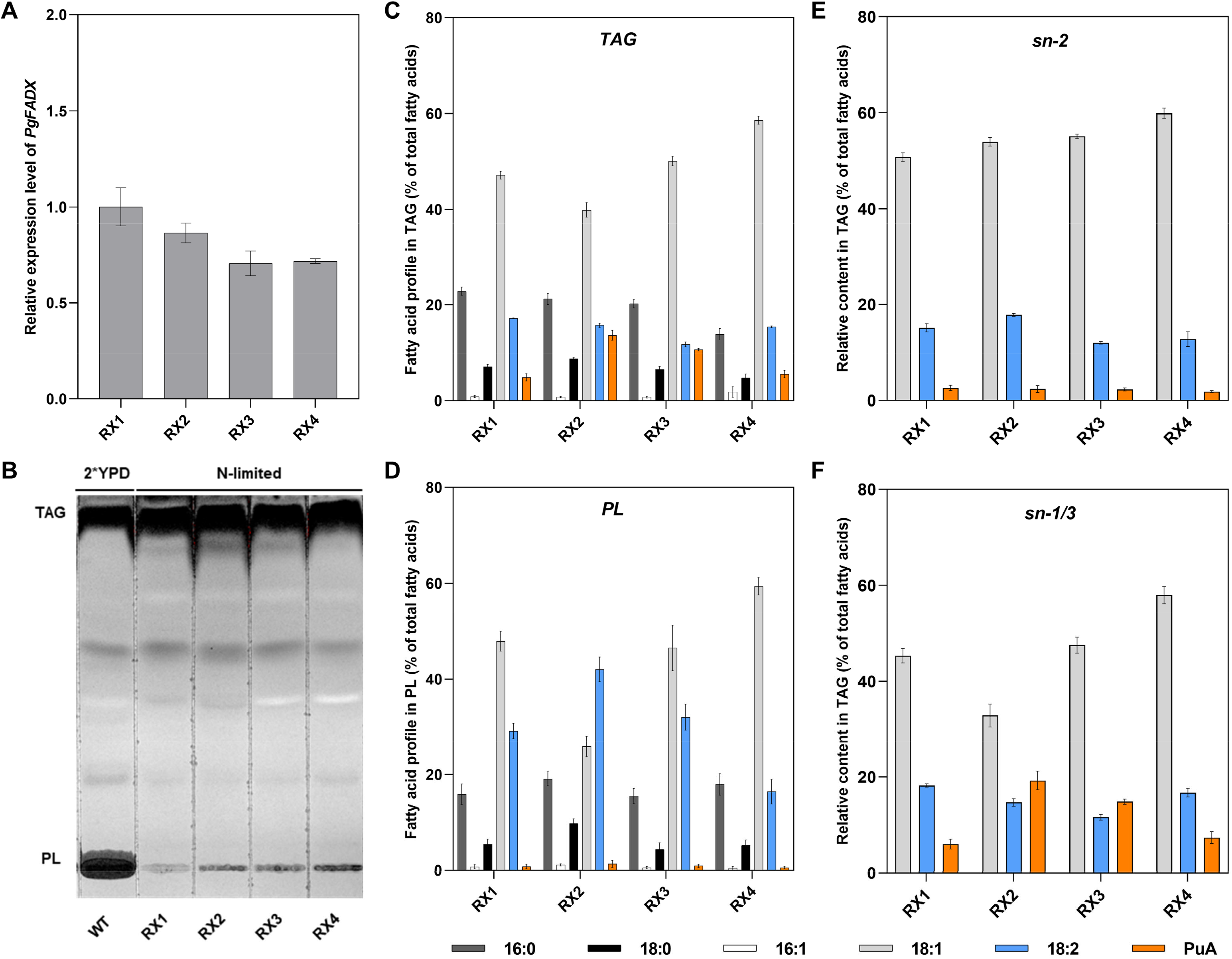
Analysis of *PgFADX* expression levels and PuA distribution in engineered *R. toruloides* strains. A) Relative expression of *PgFADX* gene in recombinant *R. toruloides* strains. *R. toruloides* GPD1 gene was used as the internal reference. B) TLC separation of lipids extracted from the wild-type and four engineered strains. C) Fatty acid composition of TAG. D) Fatty acid composition of PL. E) Fatty acid composition at the *sn-2* position of TAG. F) Fatty acid composition at the *sn-1/3* positions of TAG. Data represent mean ± SD of triplicates.

When coexpressing *PgFAD2* with *PgFADX* in *S. cerevisiae*, the LA content was increased to over 6%, whereas the PuA content was only 0.3% of total fatty acids, indicating that LA was not effectively converted to PuA (Wang et al., 2021). In the metabolic-engineered obese *Y. lipolytica* with *PgFADX* overexpression, the level of LA (2.9%) was also much higher than the PuA level (0.5%) (Urbanikova et al., 2023). In this study, the level of LA and PuA obtained in the recombinant *R. toruloides* strain RX2 was relatively more balanced, which, together with its high oleagincity and enhanced PuA content, led to a high PuA titer of 451.6 mg/L in the flask culture.

Although previous studies demonstrated that both DGAT2 and PDCT could improve the production of unusual fatty acids in transgenic hosts (Burgal et al., 2008; Yu et al., 2019), the different PuA content in RX3 and RX4 indicated the varied performance of PgDGAT2 and PgPDCT in transgenic *R. toruloides*. Indeed, many plant-derived DGAT2s have distinct acyl-CoA and DAG substrate preferences, which considerably contribute to the accumulation of unusual fatty acids (Burgal et al., 2008; Shockey et al., 2006). Consistently, *PgFADX*- *PgDGAT2* coexpression in *R. toruloides* RX3 also substantially increased PuA content compared with RX1. In contrast, although PDCT activity was proposed to play an important role in regulating unusual fatty acid exchange between DAG and PL pool, which facilitates the enrichment of unusual fatty acid in TAG (Demski et al., 2022; Yu et al., 2019), the *PgFADX*-*PgPDCT* coexpression in *R. toruloides* RX4 did not substantially enhanced PuA production (Fig. 3F). This result could be particularly due to the use of nitrogen-limited culture condition, in which the nutritional stress already limited the synthesis of PL for cell membrane proliferation, and concentrated *Rhodosporidium*’s native lipid metabolism towards TAG biosynthesis (Patel et al., 2017).

### 3.3 Distribution of punicic acid in polar and neutral lipids extracted from recombinant R. toruloides

To further elucidate the PuA distribution among different lipid classes, total lipids extracted from strains RX1, RX2, RX3 and RX4 cultured in nitrogen-limited conditions, as well as wild-type *R. toruloides* cultured in the 2*YPD medium were separated by TLC (Fig. 4B). Under the nitrogen-limited condition, all recombinant *R. toruloides* produced a higher amount of neutral lipid compared to PL. In terms of fatty acid distribution, 4.9% PuA, 17% LA, and 47% oleic acid were found in the TAG isolated from strain RX1 (Fig. 4C), while PuA made up just 0.8% of the total fatty acids in the PL fraction of RX1 lipid (Fig. 4D). With *PgFADX-PgFAD2* coexpression, strain RX2 accumulated 13.6% PuA in TAG, representing a 1.78-fold increase compared to strain RX1 (Fig. 4C). Meanwhile, the levels of LA and oleic acid were reduced by 9% and 15%, respectively. The PL fraction separated from strain RX2 lipid mainly comprises 1.4% PuA, 32% LA, and 46% oleic acid. Since both PgFAD2 and PgFADX use C18:1-PC as substrate, the amount of oleic acid in RX2’s PL was considerably reduced by 46% compared to RX1. Albeit at a lower level relative to RX2, strains RX3 and RX4 also accumulated 10.7% and 5.5% PuA in TAG, respectively, indicating a 1.2- and 0.13- fold increase compared to RX1.

In the seed lipid of pomegranate, PuA made up 60% of the fatty acids in TAG and merely 0.8% of the fatty acids in PC (Mietkiewska et al., 2014). In contrast, the distribution of unusual fatty acids between polar lipids and TAGs significantly varies in the transgenic hosts (Mietkiewska et al., 2014; Wang et al., 2021; Xu et al., 2020). In PuA-producing transgenic *Arabidopsis*, the PuA content of TAG was only 6.6%, whereas PC contains up to 12.5% of total fatty acids as PuA (Mietkiewska et al., 2014). In recombinant PuA-accumulating *S. cerevisiae,* PuA accounted for 4.8% of total fatty acids in PL, which was 1.3-fold higher than its content in TAG (Wang et al., 2021). However, in this study, recombinant *R. toruloides* led to significantly higher PuA content in the TAG than in PL. A similar observation was also reported in eicosapentaenoic acid (EPA)-producing *Y. lipolytica* (Xue et al., 2013). A possible explanation is that under nitrogen-limited conditions, TAG synthesis in oleaginous yeast underwent a significant upregulation, channelling more fatty acid towards TAG assembly instead of being utilized for membrane expansion and PL synthesis.

The positional analysis of TAG was also conducted for all four strains (Fig. 4E and 4F). Results revealed that PuA preferentially attached to the *sn-1/3* position on TAG in all four strains, with the highest percentage (19.3%) in RX2 (Fig. 4F). A similar result was found in a prior study, in which metabolic-engineered EPA-producing *Y. lipolytica* preferably concentrated EPA at the *sn-1/3* positions of TAG (Xue et al., 2013). The rearrangement of EPA across the glycerol backbone suggested significant lipid remodelling between the PL and TAG fractions in *Y. lipolytica* (Xue et al., 2013), a process that might similarly occur in *R. toruloides*. Moreover, the results also indicated that a further rationally designed metabolic engineering strategy could be developed to target the improvement of the PuA content at the *sn-2* position of TAG. Without reducing its content at the *sn-1/3* positions, such a strategy may further enrich the PuA content in the TAG fraction.

The presence of phospholipids in crude oils was sometimes considered to negatively affect the oil’s appearance and flavour (Liu et al., 2023). Since the oxidation of phospholipids has a substantial influence on the stability, shelf life, and quality of food oils, in food lipid production, degumming is often required to remove phospholipids from crude oils (Li et al., 2023). In the case of PuA-containing oils produced by heterologous hosts, removing polar lipids may result in product loss, especially if a larger ratio of PuA is retained in the polar lipid fraction rather than the neutral lipid fraction. From the biological perspective, the buildup of unusual fatty acids in PL may also lead to fitness costs (Yazawa et al., 2013; Yu et al., 2019). In this regard, the ability of engineered *R. toruloides* to accumulate a high amount of PuA in TAG under nitrogen-limited conditions makes it valuable for industrial purposes.

### 3.4 Converting wood hydrolysate into punicic acid-containing single-cell oil

*R. toruloides* has a broad substrate range and is notable for its inherent ability to metabolize pentose sugars, attracting interest for its potential in converting lignocellulosic biomass into valuable bioproducts (Sunder et al., 2023). Moreover, *R. toruloides* has shown strong growth in the presence of various inhibitors that are typically found in pretreated agricultural wastes, demonstrating its tolerance to stressful conditions compared to other yeasts (Zhang et al., 2022; Sunder et al., 2023; Kumar and Sharma, 2017). In this regard, we examined the capability of the best engineered *R. toruloides* strain RX2 in converting lignocellulose feedstock into PuA-containing single-cell oil. We first generated wood pulp hydrolysate and analyzed its sugar composition. Enzymatic hydrolysis of wood pulp yielded a liquid wood hydrolysate containing 62 mg/mL glucose and 16 mg/mL xylose, which was differentially diluted and used to replace the glucose in the nitrogen-limited medium.

As shown in Fig.5A, RX2 could successfully convert wood hydrolysate to PuA containing single-cell oil. When cultivated in 80% wood hydrolysate, RX2 accumulated 5.5% of total fatty acid as PuA. A slight increase in PuA content and titer was observed when using a higher concentration of wood hydrolysate and a higher initial OD_600_. Subsequently, a 10-day cultivation in 100% wood hydrolysate was conducted (Fig. 5B). By the end of the cultivation, the strain accumulated 6.4% of its fatty acids as PuA, and the PuA titer reached 310 mg/L in shake flask condition. The result demonstrated the potential of *R. toruloides* in producing PuA, and likely other value-added unusual fatty acids from low-cost feedstock, which is particularly valuable in sustainable bioindustry.

**Fig. 5.**
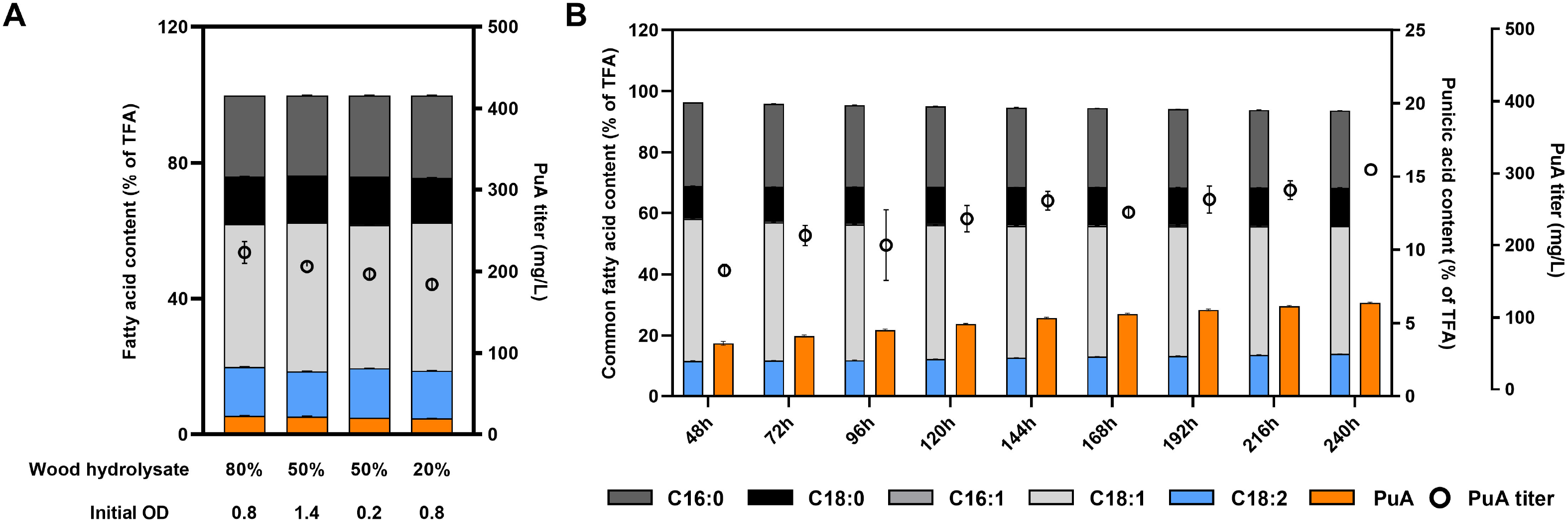
Converting wood hydrolysate into PuA-containing single-cell oil by strain RX2. A) Examining the influence of wood hydrolysate concentration and initial OD_600_ on PuA production. B) Fatty acid composition and PuA titer of *R. toruloides* strain RX2 cultured in 100% wood hydrolysate from 48h to 240h. Data represent mean ± SD of triplicates.

Lignocellulose feedstock, along with many other renewable feedstocks, is frequently regarded as a suitable substrate for microbial cell factories due to its availability and renewable nature (Sunder et al., 2023; Ling et al., 2014). The lignocellulosic biomass can be obtained from agriculture and forestry waste, and unlike starch or sugar, the abundance of lignocellulose feedstock ensures a long-term and secure supply of substrates for industrial bioprocesses without competing with food sources (Fatma et al., 2018; Kumar and Sharma, 2017). To transform lignocellulosic substrates into bioproducts, a microorganism must be capable of utilizing both pentose and hexose sugars. However, not all industrially significant strains have this capability. In the yeast *S. cerevisiae*, the introduction of heterologous xylose reductase/xylitol dehydrogenase pathway or xylose isomerase pathway was needed to enable the use of xylose as the carbon source (Gao et al., 2023). Although oleaginous yeast *Y. lipolytica* has emerged as a promising host for lipid and bioproduct synthesis, it is also unable to grow with xylose as the sole carbon source (Ledesma-Amaro et al., 2016; Zhao et al., 2015). The co-expression of xylose reductase and xylitol dehydrogenase from *Scheffersomyces stipitis* coupled with overexpression of the endogenous xylulokinase are necessary to permit normal growth of engineered *Y. lipolytica* on xylose (Ledesma-Amaro et al., 2016).

On the contrary, oleaginous *R. toruloides* can naturally assimilate pentose sugars, which is a great advantage in serving as a platform for converting low-value lignocellulose feedstock to high-value unusual fatty acid via metabolic engineering. Moreover, previous studies indicated that *R. toruloides*’ ability to convert lignocellulosic feedstock into bioproducts could be improved through the enhancement of xylose assimilation and the adaptive evolution of the engineered strain (Coradetti et al., 2023; Díaz et al., 2018; Z. Liu et al., 2021). *R. toruloides* has an unusual xylose metabolism featuring the reduction to D-arabitol, oxidation to D-ribulose, and phosphorylation to ribulose 5-phosphate (Adamczyk et al., 2023). By overexpressing a putative transcription factor (RTO4_12978, Pnt1) that acts as a major regulator of pentose metabolism, the expression of enzymes involved in xylose catabolism was increased and the specific growth rate was improved significantly in cultures on xylose (Coradetti et al., 2023). Furthermore, the adaptation of *R. toruloides* through evolutionary strategies has also led to the development of strains with increased tolerance to the major inhibitors present in lignocellulosic hydrolysates (Díaz et al., 2018; Z. Liu et al., 2021). In future, through additional adaptation of PuA-producing *R. toruloides* in lignocellulosic medium and metabolic engineering of the xylose assimilation pathway, both the cell growth and the production of the PuA could be further enhanced.

## 4. Conclusions

In this study, the non-conventional yeast *R. toruloides* was engineered for the first time to produce high-value pomegranate-derived PuA. Using glucose as the substrate, the engineered strain with *PgFADX* and *PgFAD2* coexpression accumulated 12% of total lipids as PuA with a titer of 451.6 mg/L. Utilizing wood hydrolysate as the feedstock, this strain accumulated 6.4% PuA with a titer of 310 mg/L in the shake flask culture. Taken together, the results demonstrated that *R. toruloides* could serve as an ideal metabolic engineering platform for converting agricultural and forestry waste to plant-derived high-value conjugated fatty acid.

## Acknowledgements

This work was supported by the Natural Sciences and Engineering Research Council of Canada (NSERC) Discovery Grant (D.C.B.; G.C.), NSERC Alliance Grant (G.C.), Canada Research Chairs Program (G.C.), Alberta Innovates (G.C.), Results Driven Agriculture Research (G.C.), Canada Poultry Research Council (G.C.), Canada Foundation for Innovation-John R. Evans Leaders Fund (Project number 41867), Research Capacity Program of Alberta (RCP-22-023-SEG), and Cargill/Diamond V (G.C.). We also acknowledge the financial support to students including Future Energy Systems of the University of Alberta (D.Z.H,) and Alberta Innovates Graduate Student Scholarship (J.W.)

## Data Availability

All data have been included in tables and figures.

## Declaration of interests

A provisional application has been filed (US 63/636,184), and J.W. and G.C. are listed as inventors.

