## Supplementary information for "Engineering *Rhodosporidium toruloides* for sustainable production of value-added punicic acid from glucose and wood residues"

**Supplementary Table 1** Plasmids and qPCR primers used in this study

| **Name** | **Description** | **Source** |
| --- | --- | --- |
| **Plasmids** |  |  |
| pRT-FADX | Kan^R^/ *RtCAR2-left arm*/ *P_GPD1_-codon-optimized BLE*^R^*- T_35s_* / *P_TEF1_- PgFADX-T_NOS_*/ *RtCAR2-right arm* | This study |
| pRT-FAD2-FADX | Kan^R^/ *RtCAR2-left arm*/ *P_GPD1_-codon-optimized BLE*^R^*- T_35s_*/ *P_TEF1_- PgFAD2-T_NOS_* / *P_TEF1_- PgFADX-T_NOS_*/ *RtCAR2-right arm* | This study |
| pRT-DGAT2-FADX | Kan^R^/ *RtCAR2-left arm*/ *P_GPD1_-codon-optimized BLE*^R^*- T_35s_*/ *P_TEF1_- PgDGAT2-T_NOS_* / *P_TEF1_- PgFADX-T_NOS_*/ *RtCAR2-right arm* | This study |
| pRT-PDCT-FADX | Kan^R^/ *RtCAR2-left arm*/ *P_GPD1_-codon-optimized BLE*^R^*- T_35s_*/ *P_TEF1_- PgPDCT-T_NOS_* / *P_TEF1_- PgFADX-T_NOS_*/ *RtCAR2-right arm* | This study |
| **qPCR primers** |  |  |
| qRt-GPD1-F | GTACGACCAGATCAAGCAGAC | This study |
| qRt-GPD1-R | ACAAAGTCGGTTGAGACGAG | This study |
| qRT-FADX-F1 | ACGGCTTTGCATCACACTAC | This study |
| qRT-FADX-R1 | GAGGCGGTAGAGGATGTAGTAG | This study |

**Supplementary Table 2** Codon optimized sequences used in this study

| **Name** | **Sequence** |
| --- | --- |
| PgFADX-OPTIMIZED-RT | ATGGGCGCCGACGGCACCATGTCGCCGGTCCTCACCAAGCGCCGCCCGGACCAGGAGATCAACAAGCTCGACATCAAGCCGAACCACGAGGTCGACATCGCCCGCCGCGCACCGCATTCGAAGCCGCCGTTCACCCTCTCGGACCTCCGCTCGGCCATCCCGCCGCACTGCTTCCATCGCTCACTCCTCATGTCGTCGTCGTACCTCATCCGCGACTTCGCCCTCGCCTTCCTCTTCTACCACTCGGCCGTCACCTACATCCCGCTCCTCCCGAAGCCGCTCGCCTGCATGGCATGGCCGGTCTACTGGTTCCTCCAGGGCTCGAACATGCTCGGCATCTGGGTCATCGCCCACGAGTGCGGCCACCAGGCCTTCTCGAACTACGGCTGGGTCAACGACGCCGTCGGCTTCTTCCTCCACACCTCGCTCCTCGTCCCGTACTTCCCGTTCAAGTACTCGCACCGCCGCCACCACTCGAACACCAACTCGGTCGAGCACGACGAGGTCTTCGTCCCGCGCCACAAGGACGGCGTCCAGTGGTACTACCGCTTCTTCAACAACACCCCGGGCCGCGTCCTCACACTCACCCTCACCCTCCTCGTCGGCTGGCCGTCGTACCTCGCCTTCAACGCCTCGGGCCGCCCGTACGACGGCTTTGCATCACACTACAACCCGAACGCCCAGATCTTCAACCTCCGCGAGCGCTTCTGGGTCCACGTCTCGAACATCGGCATCCTCGCCATCTACTACATCCTCTACCGCCTCGCCACCACCAAGGGCCTCCCGTGGCTCCTCTCGATCTACGGCGTCCCGGTCCTCATCCTCAACGCCTTCGTCGTCCTCATCACCTTCCTCCAGCACTCGCACCCGGCCCTCCCGCACTACAACTCGGACGAGTGGGACTGGCTCCGCGGCGCACTCGCCACCGTCGACCGAGACTACGGCTTCCTCAACGAGGTCTTCCACGACATCACCGACACCCACGTCATCCACCACCTCTTCCCGACCATGCCGCACTACAACGCCAAGGAGGCCACCGTCTCGATCCGCCCGATCCTCAAGGACTACTACAAGTTCGACCGCACCCCGATCTGGCGCGCCCTCTGGCGCGAAGCCAAGGAGTGCCTCTACGTCGAGGCCGACGGCACCGGCTCGAAGGGCGTCCTCTGGTTCAAGTCGAAGTTCTAG |
| PgFAD2-OPTIMIZED-RT | ATGGGCGCCGGCGGCCGAATGACCGTCCCGAACAAGTGGGAGGGCGAGGGCGACGAGAAGTCGCAGAAGCCGGTCCAGCGCGTCCCGTCGGCCAAACCACCATTCACCCTCTCGGAGATCAAGAAGGCCATCCCGCCGCACTGCTTCAAGCGCTCGCTCCTCAAGTCGTTCTCGTACGTCCTCTACGACCTCACCCTCGTCGCCATCTTCTACTACGTCGCCACCACCTACATCGACGCCCTCCCGGGCCCACTACGCTACGCCGCATGGCCAGTCTACTGGGCCCTCCAGGGCTGCGTCCTCACCGGCGTATGGGTCATCGCCCACGAGTGTGGACACCACGCCTTCTCGGACTACCAGTGGGTCGACGACTGCGTCGGCCTCGTCCTCCACTCGGCCCTCCTCGTCCCGTACTTCTCGTGGAAGTACTCGCACCGCCGCCACCACTCGAACACCGGCTCGCTCGAGCGCGACGAGGTCTTCGTCCCGAAGCCGAAGTCGAAGATGCCGTGGTTCTCGAAGTACCTCAACAACCCGCCGGGCCGCGTCATGACCCTCATCGTCACCCTCACCCTCGGCTGGCCGCTCTACCTCGCCCTCAACGTCTCGGGCCGCCCGTACGACCGCTTTGCATGCCACTTCGACCCGTACGGCCCGATCTACACCGACCGCGAGCGCCTCCAGATCTACATCTCGGACGTCGGCATCATGGCCGCCACCTACACCCTCTACAAGATCGCCGCCGCCCGAGGACTAGCCTGGCTCGTCTGCGTCTACGGCGTCCCGCTCCTCATCGTCAACGCCTTCCTCGTCACCATCACCTACCTCCAGCACACCCACCCGGCCCTCCCGCACTACGACTCGTCGGAGTGGGACTGGCTCCGCGGCGCACTAGCCACCGCCGATCGAGACTACGGCATCCTCAACAAGGTCTTCCACAACATCACCGACACCCACGTCGCCCACCACCTCTTCTCGACCATGCCGCACTACCACGCCATGGAGGCCACCAAGGCCATCAAGCCGATCCTCGGCGACTACTACCAGTTCGACGGCACCCCGGTCTACAAGGCCATGTGGCGCGAGGCCCGCGAGTGTCTCTACGTCGAGCCGGACGACGGCGCCAACTCGAAGGGCGTCTTCTGGTACAAGAAGAACCTCTAG |
| PgDGAT2-OPTIMIZED-RT | ATGGGCGAGGAGGCCTCGGTCAAGCTCGGCGAGGACCAGCAGCGCGAGGTCTTCACCGGCCGCAAGGAGTCGCCGTCGCCGGTCACCTTCCACGCCCTCCTCGCCCTCGCCATCTGGGTCGGCACCATCCACTTCTTCGGCTTCCTCGTCTCGGTCTCGCTCCTCGTCCTCCCGCTCTCGAAGGCCCTCCTCGTCTTCGGCCTCCTCGGCGCCCTCGTCGTCATCCCGGCCGACGACCGCTCGAAGTTCGGCGAGCGCGTCACCCGCTACATCCTCAAGCACGCCTGCCCGTACTTCCCGATGACCCTCCACGCCGAGGAGTTCGGCTGCATCGACCCGAACCGCGCCTACGTCTTCGGCTACGAGCCGCACTCGGTCATGCCGGTCGGCACCGTCGCCCTCGCCCAGCTCTACGGCCTCATCACCATCCCGAAGCTCAAGGTCCTCGCCTCGACCGTCGTCTTCCGCACCCCGTTCATCCGCCACGTCTGGACCTGGATGGGCCTCACCCCGGCCACCCGCAAGAACTTCATCTCGCTCCTCGAGTCGGGCTACTCGTGCATCGTCGTCCCGGGCGGCGTCCAGGAGACCTTCTACATGGAGCACGGCTTCGAGGTCGTCTTCCTCAAGAAGCGCCGCGGCTTCGTCCGCATCGCCCTCGAGACCGGCTGCCCGCTCGTCCCGGTCTTCTGCTTCGGCCAGTCGCAGCTCTACAAGTGGTGGAAGCCGTCGTGGGGCCTCTTCCTCAAGATCTGCCGCGTCGTCAAGTTCACCCCGATGTTCTTCTGGGGCATGCTCGGCTCGCCGCTCCCGTTCCGCCACCCGCTCCACATCGTCGTCGGCAAGCCGATCGAGGTCAAGCGCACCCCGAACCCGACCGCCGAGGAGGTCGACGAGCTCCACAAGCAGTACGTCGAGGCCCTCCGCGACCTCTTCGAGCGCCACAAGGCCCAGGTCGGCCACGAGGACCTCGTCCTCAAGATCCTCTAG |
| PgPDCT-OPTIMIZED-RT | ATGAACGGCGCCAAGTCGACCACCGCCACCATCACCCGCCGCCGAGACCCACGCTCGCCATCGAACGGCCTCGTCCTCGACCCGGTCGCAGGAATGGCCAACGGAAAGCGCGCAGTCGTCAACGGCGGCTACGGAGACTACAACAAGGCCAAGGCCGCCGTCGCCTTCATGCGCTGGACCCGCGACGACGTCTTCAACCTAGCACGCTACCACCGCCTCCCGTGCCTCTTCGCAGCCGGCCTCCTCTTCTTCATGGGCGTCGAGTACACCCTCCTCATGGTCCCGGACGACCTCCCGCCGTTCGACCTCGGCTTCGTCGCCACCCGCTCGCTCCACCGAGTCCTATCGTCGTCGTCGGAGCTCAACACCATCCTCGCCGCCCTCAACACCGTCTTCGTCGGCATGCAGACCGCCTACATCCTCTGGGCCTGGCTCATCGAGGGCCGCCCACGAGCCACCATCTCGGCACTCTTCATGTTCACCTGCCGCGGCATCCTCGGCTACTCGACCCAGCTCCCGCTCCCGCAGGGCTTCCTCGGCTCAGGCGTCGACTTCCCGGTCGGCAACGTCTCGTTCTTCCTCTTCTTCTCGGGCCACGTCGCCGGCTCAGTCATCGCCTCGCTCGACATGCGCCGCATGAAGCGCTGGGAGCTCGCCTGGACCTTCGACGTCCTCAACGTCCTCCAGGCCGTCCGCCTCCTAGGCACACGAGGCCACTACACCATCGACCTCGCCGTCGGCCTCGGAGCAGGCATCCTCTTCGACTCGCTCGCCGGCAAGTACGAGGAGTCGCACAAGATGCGCAAGGGCATCATCCACCACGTCAACGGCATGAACGGCTCGAAGGAGGGCCCGATGATCTAG |
| TEF1 PROMOTER | CGCGAAGCGGTAGAAGCAATGAAGCGAGGCGAGAGCGAGAGAGGCAGGGCTTCAGCCATGTCCAGCTGATCGGCTGTAACGTCGCGCCGGGCCAGTCTGTTGAATTTGTTGCGTCGCCTGAGCGTAATAGAAGTGCAGTAGTCTACTCCGCATGCCGAGAACGTCGAAGAGCGCGAAGTAGGGAGTCGAGGGAAGCGAGGGTGGCAAACACAGCAACGACAAGCGGTTCCGCTTCGCTCAAAAGCTCGTTGACGTTGTTTTGACGTTTTGAAGACAGTACAACAGCAGCAAGAGGCGTGCGAAGCGTTGGTGGCGAGAGCAGCGACAAGGAGGGAGGAATGAGGGAGTGGTGGCGAGGGCTCGCAAACGGGCGTACGCCTCGAATGGAGACGTGCGAGTCGTTCTTCGACGTCCGAGGGATGCCGAGCGCCGAGACGGAGCACGCAACGAGCGAGAGGAGAGCAGCCGCGCAAGGTGATTCGAGTGGCGCAAGCGGAGGACGACGAGGAGACGGACGAGGGAGGAGGAGGGATGGCGAGCGAGCATCGGACGGCGGGGCGCGAGAGACGGCGTGAGGAGCCGGGTGTGGAGAGTTTGAGGAGGCGCGGGATGCGAAGTGGCTGGGTGTGCGGAGTGAGCGGTGGCAAAGAGCGCACTTAGAGTCTAGAGCGAGGCAGTAGTAGTAGAGCTGTATGAATGAATACAAAGTGTGAATACAACAGTTTGTAATGCGATTCTGAGCTTGGACGTGTGCGCGCGAGAGGGCGACTTGCAAGCCAGCGCCCGCTCGCTCTTCTTCCTTCTGCACCTCGCGTCAACCCTCGCATCTCACACCTACACTCGCATTCAAAGTGCGTACACTCTCCCACGACACACGGGGACGGCGCACACCACCGCGCGTCGCTTGAACGGCGTCGCCACTTCGAGCCGTCACTGACTTCGTCCTCGTCCTCCCTCCTCTACTCTCTTGTACTGTActgtgtactgggggggatag |
| RtGPD1 promoter | TCTTCAGACGGCTTGTTCTCTCCTGCTCTGGTGGGCTGGCCTGACATGTAATGTGCTCCGCCGCAAGTCCGTCGTCGGTCTCAATTCGACGTTGAAAGGGCATAGCGCAAGGAAGAACCCTCTGCGGACATGCAGAATTACTGGCTCGCCTGCTCCTTCGTCTACTGGAATAAGTCCTGTCTCGTTAAAGCCCCAACGTCGTTTTTCGACGTTTGTAAGGCGCAAGAGGTGCTATGGGCTACGCAGGAAGCTGAGAGGACATAGAAGTCGGGGGAGGAACGGCGCAGAGCGGCAGTTGCGGAAGCATGAGGAAAGCGAGACGGTCCAGCATCTGCAGCGCCAATCCGCAATCTCCTGGTTGAGCCTGCACCGGAAGCGTCGGAACAGTATGCGCAGAGTCGAACGCAAGTAAGAAAGACGCACCCTCACACTCGCTTACTTCGAGCCATACAACGGATCAAAGCTGCGCGTATCTCGGCTTGTAAGGGCCGGAAAGCAACCTCGGAGATGGACACGTCACATCACCAACTTATCGATCTCGGCCGTCGACGTCGCAGAGAGGGCGAGAGAAGCGGTGAAGGAGGGAAACAACCCCTCGAGAGCATGATCCGACCGAATCTGCAGCGCAGGAAGCCGTTACAAGCCCGCCTCGAGCGCAGGTCGGGTCCAGCCGGGGGACGAAACGCGCGAGGCTGATTCGTGAGCGAAGGAAGCCGCATCGACAAGTTCGCTCCCCTTTGCCCTCTTTCCCATCACCCGTTCTCGCCTTACCCGCTCAGAACAACACCAGATCACTCACA |
| BLE-OPTIMZED-RT | ATGGCCAAGCTCACCTCGGCCGTCCCGGTCCTCACCGCCCGAGACGTCGCAGGAGCAGTCGAGTTCTGGACCGACCGCCTCGGCTTCTCGCGCGACTTCGTCGAGGACGACTTCGCCGGCGTCGTCCGCGACGACGTCACCCTCTTCATCTCGGCCGTCCAGGACCAGGTCGTCCCGGACAACACATTAGCATGGGTCTGGGTCCGAGGCCTAGACGAGCTCTACGCCGAGTGGTCGGAGGTCGTCTCGACCAACTTCCGCGACGCCTCGGGACCAGCCATGACCGAGATCGGCGAGCAGCCGTGGGGACGCGAGTTCGCCCTCCGCGATCCGGCAGGCAACTGCGTCCACTTCGTAGCCGAGGAGCAGGACTAG |
| 35S terminator | acgctgaaatcaccagtctctctctacaaatctatctctctctattttctccataaataatgtgtgagtagtttcccgataagggaaattagggttcttatagggtttcgctcatgtgttgagcatataagaaacccttagtatgtatttgtatttgtaaaatacttctatcaataaaatttctaattcctaaaaccaaAATCCAGTACTAAAATCCAGA |
| NOS terminator | gatcgttcaaacatttggcaataaagtttcttaagattgaatcctgttgccggtcttgcgatgattatcatataatttctgttgaattacgttaagcatgtaataattaacatgtaatgcatgacgttatttatgagatgggtttttatgattagagtcccgcaattatacatttaatacgcgatagaaaacaaaatatagcgcgcaaactaggataaattatcgcgcgcGGTGTCATCTATGTTACTAGATC |
